# Retinoic acid promotes in vitro development of haploid germ cells from pre-pubertal porcine spermatogenic cells

**DOI:** 10.1101/552083

**Authors:** Kun Yu, Yi Zhang, Bao-Lu Zhang, Han-Yu Wu, Su-Tian Wang, De-Ping Han, Zheng-Xing Lian, Yi-Xun Liu, Shou-Long Deng

## Abstract

Spermatogonial stem cells (SSCs) self-renew and contribute genetic information to the next generation. Inducing directional differentiation of porcine SSCs may be an important strategy in exploring the mechanisms of spermatogenesis and developing better treatment methods for male sterility. Here, we established an in vitro culture model for porcine small seminiferous tubule segments, to induce SSCs to differentiate into single-tail haploid spermatozoa. The culture model subsequently enabled spermatozoa to express the sperm-specific protein acrosin, and oocytes to develop to blastocyst stage after round spermatid injection. The addition of retinoic acid (RA) to the differentiation media promoted the efficiency of haploid differentiation. RT-PCR analysis indicated that RA stimulated the expression of Stra8 but reduced the expression of NANOS2 in spermatogonia. Genes involved in post-meiotic development, Prm1 and Tnp1, were up-regulated in the presence of RA. The addition of RAR inhibitor, BMS439, showed that RA enhanced the expression of cAMP responsive-element binding protein through RAR, and promoted the formation of round spermatids.

## Introduction

Mammalian spermatogonia originate from primordial germ cells. In rodents, type A_single_ spermatogonia (As) undergo self-renewal and proliferate into type A_paired_ spermatogonia (Ap) to initiate the process of spermatogenesis. These spermatogonia subsequently form type A_aligned_ spermatogonia (Aal) and finally turn into type A1 spermatogonia without mitosis. These type As, Ap and Aal spermatogonia are collectively called undifferentiated spermatogonia. Some genes involved in the self-replication of spermatogonial stem cells (SSCs), such as NANOS2, are essential to ensure a stable number of stem cells. NANOS2 is a conserved zinc-finger RNA-binding protein that maintains the self-replication of As and Apr spermatogonia [1]. Its continuous expression is regulated by glial cell-derived neurotrophic factor (GDNF) through the GDNF family receptor alpha 1 (GFRα1) on SSCs [2, 3]. Type A1 spermatogonia then undergo mitosis and give rise to a series of differentiating spermatogonia types (A2, A3, A4, intermediate (In) and B type) before initiating meiosis as preleptotene primary spermatocytes. These differentiated type A1-derived spermatogonia express Kit and mitotic genes, which are specifically expressed before meiosis and the stimulated by retinoic acid gene 8 (Stra8) [4]. The last phase of spermatogenesis is spermiogenesis, in which round haploid spermatids develop into mature flagellated spermatozoa. Spermatogenesis involves complex stages of cell differentiation and requires the involvement of various key factors, such as support cells, essential nutrients (amino acids, vitamins) and reproductive hormones (testosterone, follicle stimulating hormone (FSH), luteinizing hormone (LH)), as well as synergism between cytokines [5, 6], and construction of the required gene regulation network in spermatogenic cells [7]. Several culture systems have been developed to investigate the complete process of spermatogenesis in vitro [8, 9]. However, owing to limited differentiation efficiency, these in vitro models are not ideal for the practical production of functional sperm [10, 11].

Previous research reported that retinoic acid (RA) at a concentration of 10^-8^ M was sufficient for activating Stra8 and promoting the onset of meiosis [12, 13]. Retinoic acid is a metabolite derived from vitamin A [14]. When bound to its high affinity retinoic acid receptor (RAR), RA affects the RA response elements in promoters of target genes to regulate transcription. The RAR includes three isomers, RARα, RARβ and RARγ. In newborn, pubertal and adult mammalian testes, RARα is mainly located in testicular Sertoli cells. Retinoic acid receptor gamma is mainly expressed in differentiated spermatogonia. Retinoic acid regulates spermatogonia differentiation mainly through RARγ [15].

Retinoic acid deficiency leads to elevated SSC numbers in the neonatal mouse testis [16]. The differentiation of spermatogonia needs RA [17]. In mice, long term vitamin A deficiency or retinoic acid antagonist (such as WIN18446) administration will block spermatogenesis at the early undifferentiated (Aal) stage, and result in azoospermia and infertility. Replenishment of vitamin A or RA can restore fertility by inducing spermatogonial maturation from type Aal to type A1 [18]. In Sertoli cells, RA enhances the expression of Kit ligand (KL, the Kit receptor) and bone morphogenetic protein 4 (BMP4), which inhibits the expression of GDNF [19]. In undifferentiated spermatogonia, RA combines with RARγ to stimulate the expression of Kit [20] and Stra8 genes [21]. Recent studies have examined the effect of RA on inducing the differentiation of cultured SSCs in vitro [22]. The miniature pig is an ideal animal model for understanding human reproduction, with advantages including similarities between mini-pig and human anatomy, physiology and pathology, and the benefit of short estrous cycles and a large number of piglets [23, 24]. This study will build the foundation for accomplishing porcine spermatogenesis from SSCs in vitro, and ultimately contribute to a better understanding of the mechanism of RA action during the initiation of meiosis and sperm formation.

## Materials and methods

### Isolation of piglet seminiferous tubule fragments

Testis tissue was obtained from 2-month-old Chinese experimental miniature pigs. Testes were transported to the laboratory in phosphate buffered saline (PBS) supplemented with 100 mg/mL streptomycin and 100 IU/mL penicillin. After decapsulation, seminiferous tubule fragments were dissociated by modified enzymatic digestion [25, 26]. Briefly, the seminiferous tubules were incubated with an enzyme cocktail containing 0.1 mg/mL collagenase type IV and 1.0 μg/mL DNaseI at 37°C for 15 min, followed by neutralization with 10 % fetal bovine serum (FBS) (Gibco, Grand Island, NY, USA). The suspension was filtered using a 40 mesh sieve. The seminiferous tubule fragments were cultured in medium containing DMEM/F12, 20% KnockCut Serum Replacement (KSR), 2 mmol/L L-glutamine, 1% non-essential amino acids, 10 ng/mL fibroblast growth factor, 20 ng/mL GFRα1, 10 ng/mL GDNF, incubated at 37°C in 5% CO_2_/air for 3 d.

### *In vitro* differentiation of SSCs

The seminiferous tubule fragments were suspended in medium (M) containing DMEM, 5% KSR, 1% non-essential amino acids, 10 ng/mL stem cell factor, 10 ng/mL fibroblast growth factor, 25 ng/mL epidermal growth factor, 10 ng/mL insulin-like growth factor, 10 μg/mL transferrin, 2 mM L-glutamine, 0.05 IU/mL FSH, 0.05 IU/mL LH, 0.1 μmol/L testosterone and 1% penicillin-streptomycin. The temperature was then maintained at 34°C in 5% CO_2_/air. On the fifth day of culture, 1 mol/L RA was added to one group and after 48 h of incubation, the medium was replaced by normal medium for further culture. One group of media was then supplemented with 1 mol/L RA to for 96 h. To test for RA-specific actions, we added 5 μM BMS493, a pan-RAR antagonist [27]. In each culture system, half the medium was changed every 2 d. The rate of cell growth was observed.

### Quantitative real-time PCR

The prepared cells were collected to determine gene expression levels. Total RNA was extracted using Trizol reagent (Invitrogen, Carlsbad, CA, USA) according to the manufacturer's protocol. Reverse transcription-PCR was performed using a cDNA synthesis kit (Promega, Madison, WI, USA) and 2 μg of total RNA according to the manufacturer's protocol. The SSC related genes (GFRα1, PGP9.5, Plzf and NANOS2 [28]), SSC differentiation-related genes (Stra8, c-kit, RARγ, and cytochrome P450 family 26 enzymes B1 (CYP26B1)), an anti-apoptosis gene (Bcl2), genes with post-meiotic expression (transition protein 1 (Tnp1) and protamine 1 (Prm1)), and histone acetylation-related genes (Cdyl and Hdac1) were detected by RT-PCR. β-actin was used as an internal control. The primer sequences are listed in Table 1. Real-time PCR reactions were carried out with a Real Master Mix SYBR Green Kit (Tiangen, Corp, Beijing, China) using a Stratagene Mx300p (Agilent Technologies Inc, Santa Clara, CA, USA). Fold change of gene expression was calculated using the 2^-ΔΔ*ct*^ method, and was expressed as a ratio of expression levels of treated groups to the expression level of the control group.

**Table 1.**
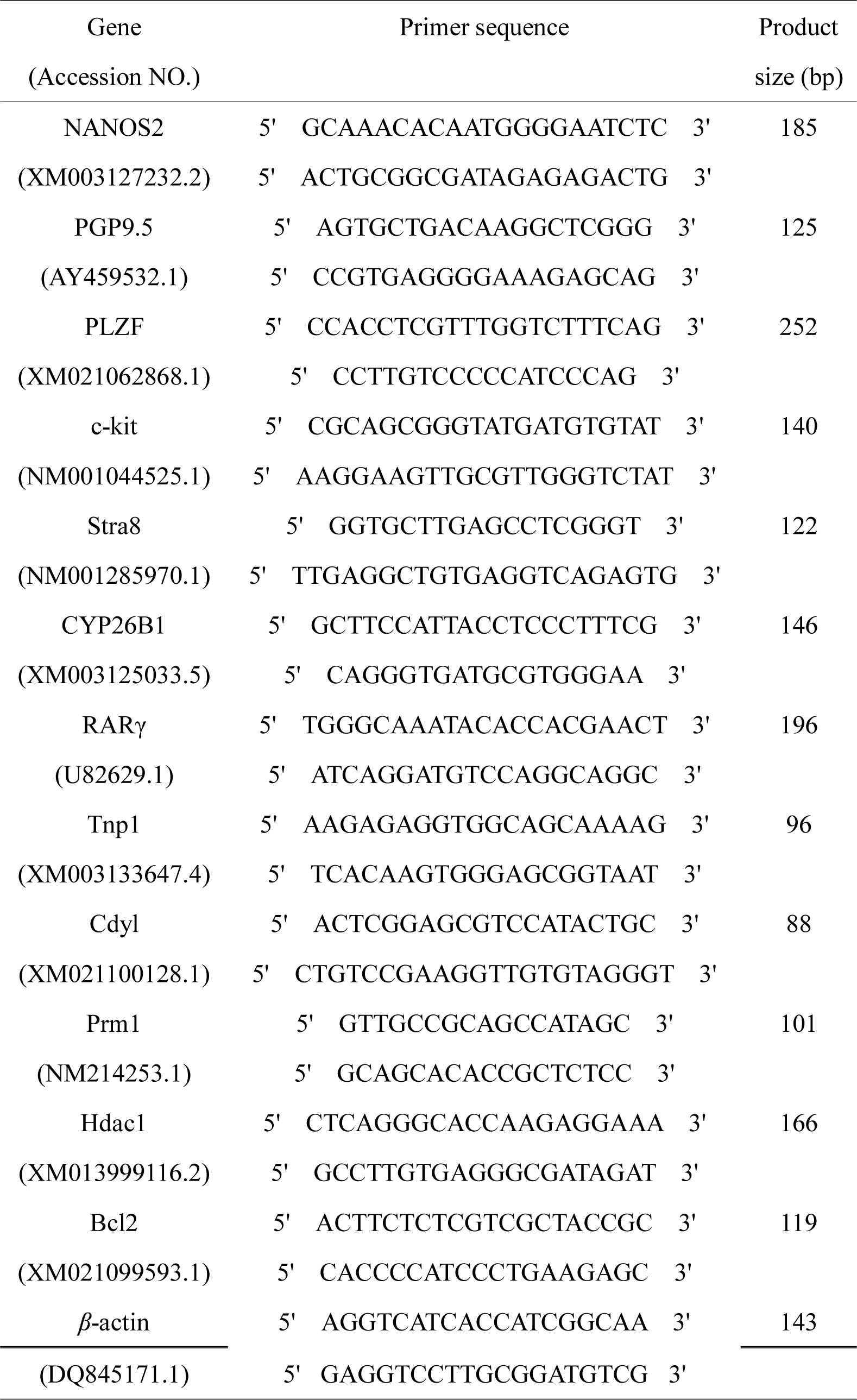
Primer sequences.

### Flow cytometric analysis

The DNA content of cells was examined by flow cytometry. *In vitro* cell suspensions adjusted to 1×10^6^ cells/mL were collected at 9 days, and sperm from a mature pig was used as a control. The cells and sperm were fixed in 70% ethanol for 4 h. After three washes in PBS, the cells were incubated at 37°C for 10 min in PBS plus 200 μg/mL RNase I and 20 μg/mL propidium iodide (PI). Cells cultured for 5 days were examined by flow cytometry for germ cells. Briefly, the cells were fixed in 70% alcohol for 2 h, then washed twice in PBS, and then re-suspended in PBS with BSA for 1 h. The cells were then incubated with anti-UCHL1 antibody (Santa Cruz Biotechnology, Santa Cruz, CA, USA, sc-25800, diluted 1:200), GFRα1 (Santa Cruz Biotechnology, sc-6157, diluted 1:200) and anti-CDH1 antibody (Santa Cruz Biotechnology, sc-1500, diluted 1:200) for 1 hour. Cells were then washed three times in PBS by centrifugation at 500×g for 5 min, and then secondary antibody was added and incubated for 45 min. The cells were then washed three times with PBS and re-suspended in 0.5 mL PBS for analysis by flow cytometry.

### Immunofluorescence analysis and ELISA

Cells were examined by immunofluorescence staining after 3 days of culture for GFRα1, a marker for SSCs [29], after 7 days of culture for Stra8 (Abcam Inc., Cambridge, MA, USA, ab49602), a marker for differentiated spermatozoa, and after 9 days of culture for acrosin (Bioss, Beijing, China, bs-5151R), a marker for spermatozoa [30]. Briefly, cells were fixed in 70% alcohol for 2 h, and then washed twice in PBS. Slides were blocked with 1% BSA for 1 h at room temperature, and primary antibody (diluted 1:200) was added to the solution and incubated for 4 h. Slides were rinsed twice and washed three times with PBS for 5 min. Secondary antibody (1:500) was incubated for 1 h at room temperature, followed by the same PBS washes, and nuclei were stained with DAPI. Cells cultured for 9 day were collected for cAMP responsive-element binding protein (CREB) detection by an ELISA kit (Hermes Criterion Biotechnology, Vancouver, Canada) following the manufacturer's protocol.

### Immunohistochemistry

Testis samples from 2-month-old pigs were fixed with 4% paraformaldehyde. The samples were cryo-embedded in OCT compound, and then cut into 7 μm thick sections and stained using hematoxylin and eosin (H&E) for histological analysis of the seminiferous tubules. The Uchl1 [31] and RARα [32] (Santa Cruz Biotechnology, sc-551) expression patterns were examined by immunohistochemistry. Briefly, after washing three times with PBS, the slides were incubated in PBS containing 1% BSA for 1 h at room temperature. Primary antibodies (diluted 1:200) were added to the solution respectively. After 4 h of incubation, the secondary antibody was applied for 1 h. Staining was visualized using a DAB substrate kit.

### Intracytoplasmic microinjections

Ovaries were collected from a slaughterhouse. Cumulus cells and cumulus-oocyte complexes (COCs) were selected and cultured in *in vitro* maturation (IVM) medium that included 10% FBS TCM-199 (Gibco), 10 ng/mL epidermal growth factor, 10% porcine follicular fluid, 10 IU/mL equine chorionic gonadotrophin, 5 IU/mL human chorionic gonadotrophin, and 2 mM glutamine. Oocytes extruding the first polar body were selected for injection of round spermatids collected from testes [33]. Spermatids less than 10 µm in diameter with single flagella were collected from the in vitro system and used for microinjection [34]. Micromanipulation was performed in TCM-199 medium supplemented with 5 μg/mL cytochalasin B, 3 mg/mL BSA, and 0.5 mM HEPES. Oocytes were activated with 5 μM ionomycin for 5 min before injection. Cell membranes of spermatids were disrupted by repeated blowing with an injection needle, and then spermatids were injected into the cytoplasm of oocytes. Intracytoplasmic injection was finished within 1 h after activation. Recovered couplets were transferred into development medium, porcine zygote medium (PZM-3), for recovery at 38°C and 5% CO_2_ for 30 min, and then activated with 10 μg/mL cycloheximide and 10 μg/mL cytochalasin B for 4 h. After activation, reconstructed embryos were cultured at 38°C in 5% CO_2_ for development, and the development of double pronuclei in reconstructed embryos was observed by lichen red staining, and blastocysts were observed on day 7.

### Statistical analyses

All experiments were repeated at least 3 times. One-way ANOVA was used to determine statistical significance with the Duncan’s test used to determine the statistical significance between the relative groups. Statistical analysis was conducted using Statistical Analysis System software (SAS Institute, Cary, NC, USA). All data were expressed as mean ± SEM. Differences were considered to be significant when *P* < 0.05.

## Results

### Identification of pig SSCs

The hematoxylin and eosin-stained sections of 2-month-old porcine testis revealed that only undifferentiated spermatogonia were present in seminiferous tubules (Figure 1A). Expression of Uchl1 was detected in porcine SSCs by immunostaining (Figure 1B). Both Sertoli cells and spermatogenic cells expressed RARα (Figure 1C). Adherent Sertoli cells had grown out from small seminiferous tubule segments when observed 3 days after plating. At this stage, some spermatogenic cells gathered around the Sertoli cells, free but close to the surface of Sertoli cells (Figure 1D). Thereafter, bridge and chain connections between cells were observed (Figure, 1E), and these cells expressed GFRα1 (Figure 1F). On day 5 of RA treatment, SSC colonies were observed (Figure 1G). Expression levels of GFRα1, PGP9.5, PLZF and NANOS2 transcripts were significantly higher on day 5 compared with day 3 of incubation (Figure 1H), and simultaneous flow cytometric analysis identified UCHL1, GFRα1 and CDH1 protein expression in the culture system (Figure 1K).

**Figure 1.**
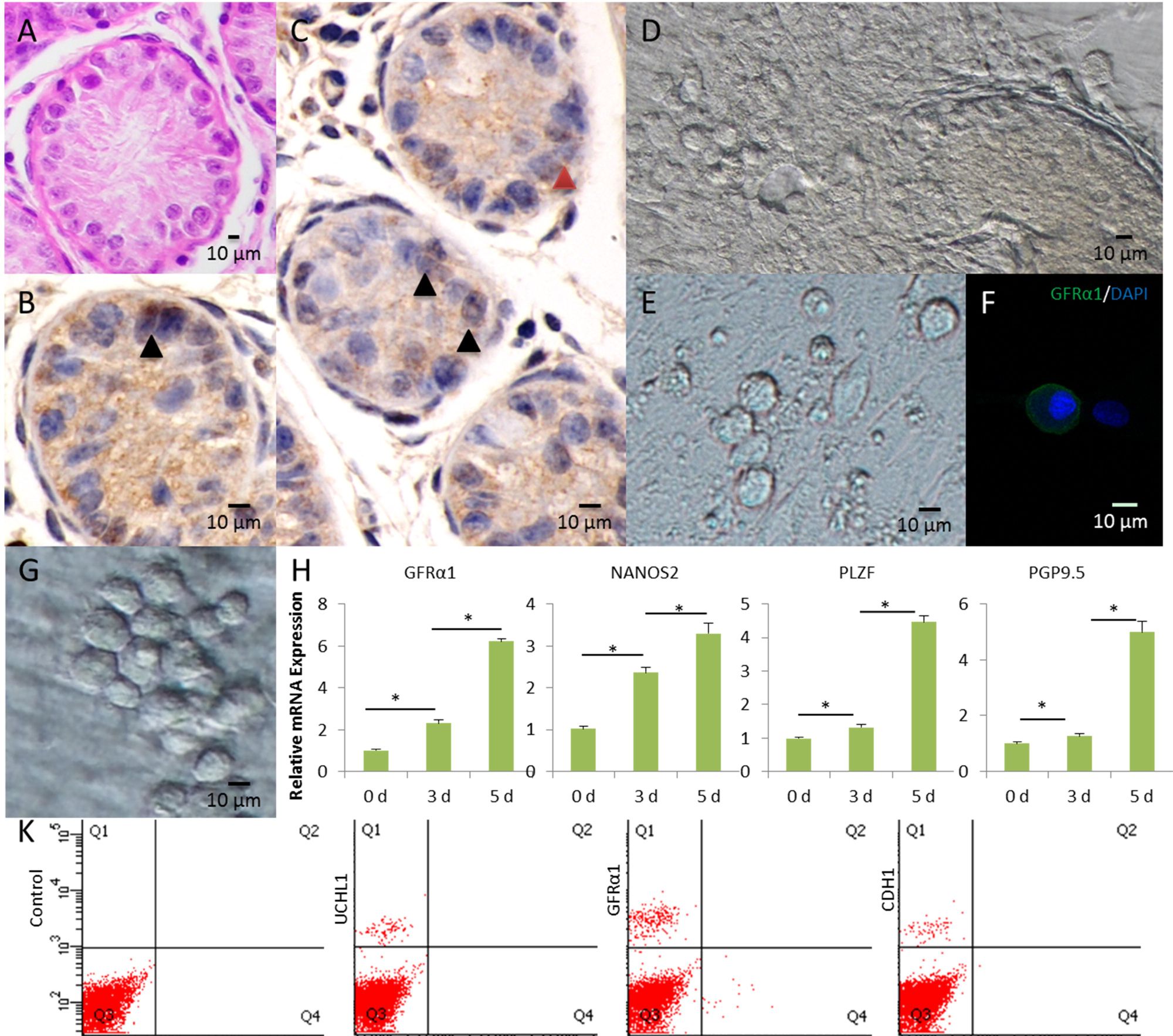
Male spermatogonial stem cell (SSC) culture. A) H&E staining of the 2-month-old pig testis. B) Immunohistochemical analysis of UCHL1 expression in the 2-month-old pig testis. Primordial germ cells are indicated by black arrows. C) The expression of RARα in the seminiferous tubules of the porcine testis. Primordial germ cells are indicated by black arrows and Sertoli cells by red arrows. D) In vitro culture of porcine seminiferous tubules. E) Bridge and chain connections between SSCs in vitro. F) Immunofluorescent analysis: GFRα1 (green), DAPI-stained nuclei (blue). G) SSC colony. H) Real-time PCR analysis of GFRα1, PGP9.5, PLZF and NANOS2 mRNA levels in the in vitro system at various times (day 3 and 5). Data are expressed as means ± SEM; \**P* < 0.05. K) Flow cytometric analysis of UCHL1, GFRα1 and CDH1on day 5 of incubation.

### RA up-regulated the expression of STRA8 in porcine SSCs in vitro

After 48 h induction of SSCs on day 5, the expression of Stra8 was localized to spermatogenic cells (Figure 2A). Expression of RARγ mRNA levels was significantly elevated in the RA group compared with the M group (*P* < 0.05). The expression of gene Stra8 and c-kit was also significantly higher in the RA group than that in M group (*P* < 0.05), indicating that RA may promote the expression of Stra8 and c-kit through its receptor (Figure 2B). Expression of NANOS2 and GFRα1 mRNA levels was reduced in the RA group compared with the before induction (*P* < 0.05). Reduced expression of PLZF mRNA was also found in the RA-treated relative to the M group (*P* < 0.05), however, there was no significant difference compared with the control group (Figure 2C), suggesting that RA induced SSCs to initiate meiosis. Additionally, decreased expression of CYP26B1 mRNA was observed in the RA group compared with the M and control groups. These results suggest that RA reduced the expression of NANOS2, GFRα1 and PLZF in spermatogonial cells, and promoted the expression of Stra8 in meiotic spermatogenic cells.

**Figure 2.**
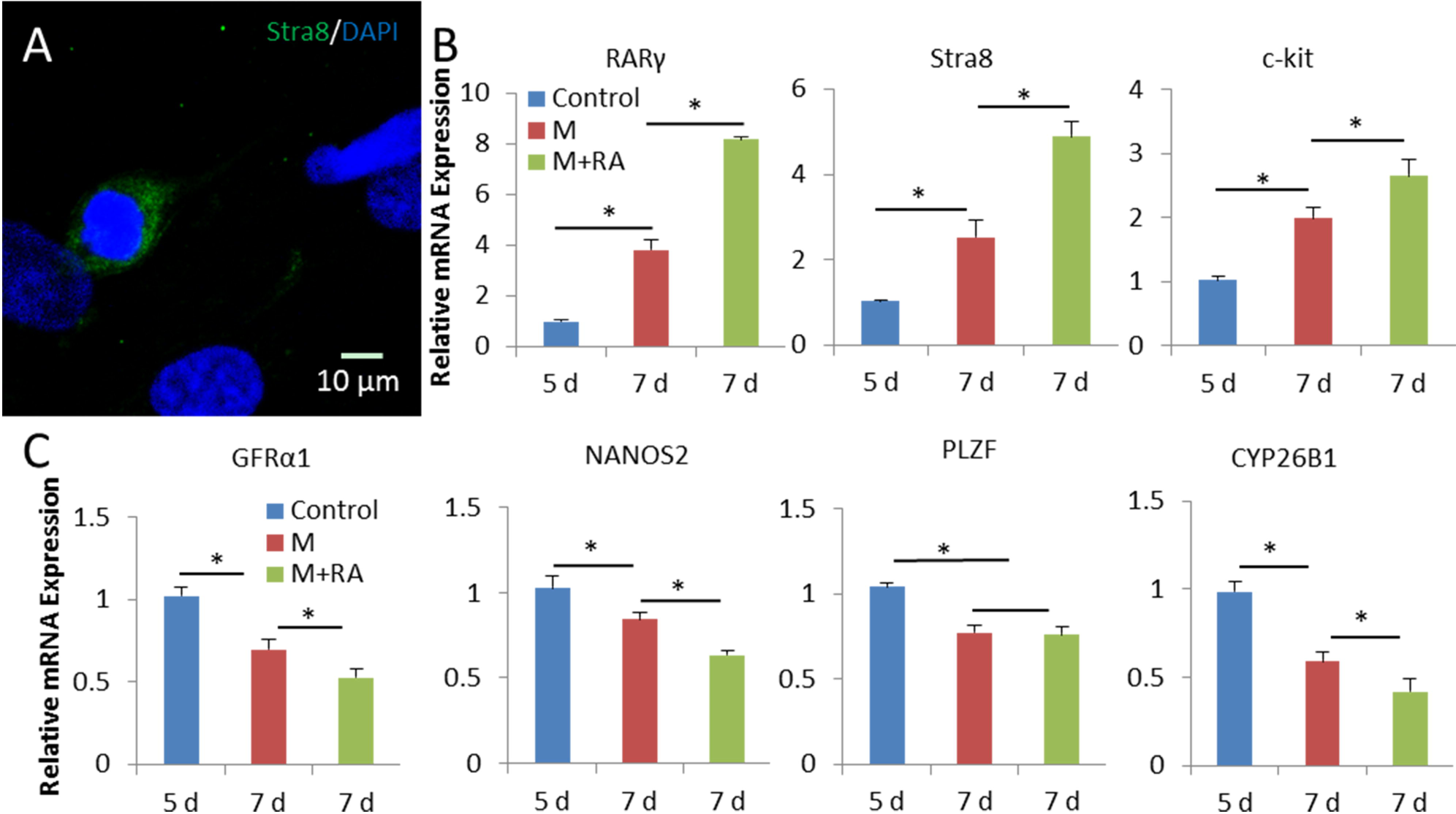
Retinoic acid up-regulated the expression of Stra8 in the porcine SSC in vitro differentiation system. A) Immunofluorescent analysis: Stra8 (green), DAPI-stained nuclei (blue). B) and C) Real-time PCR analysis of RARγ, Stra8, c-kit, GFRα1, NANOS2, PLZF and CYP26B1 mRNA levels in the in vitro system at various times (day 5 and 7). Control is the group without induction (SSCs on day 5 of incubation), M is the group that was induced to differentiate with basic medium, and M+RA is the group with RA treatment. Data are expressed as mean ± SEM; \**P* < 0.05.

### In vitro differentiation of SSCs into sperm-like cells

The differentiation of a single tail to Sa spermatid or Sd type spermatozoa (Figure 3A and B) (Supplementary movie) was observed on day 9 of incubation. Immunofluorescence showed some cells in the culture system expressed round spermatid-specific acrosin (Figure 3C). Ploidy analysis revealed that the haploid efficiency of the M + RA-96h group was 5.3 ± 0.83 % higher than that of the M + RA-48h group and M group (Figure 3D and Table 2). At the later stage of meiosis, Tnp1 and Prm1 were expressed at significantly higher levels in the M + RA-96h group than in the other groups (Figure 3E and F). There was no significant difference in histone acetylation modifying enzyme Cdyl and Hdac1 (coding histone deacetylase) gene expression within the groups (Figure 3G and H). The above results indicate that in the in vitro induction culture system of spermatogenic cells, continuous RA treatment can significantly increase the differentiation rate of haploid cell and sperm formation in vitro.

**Table 2.** Ratio of haploid spermatozoa in suspended cells 9 days after SSC differentiation.

| In vitro differentiation medium | Haploid ratio (%) |
| --- | --- |
| M | $1.7 \pm 0.32^c$ |
| M+RA-48h | $2.9 \pm 0.55^b$ |
| M+RA-96h | $5.3 \pm 0.83^a$ |

Note: <sup>a, b, c</sup> Difference among M, M+RA-48h and M+RA-96h ( $P < 0.05$ ).

**Figure 3:**
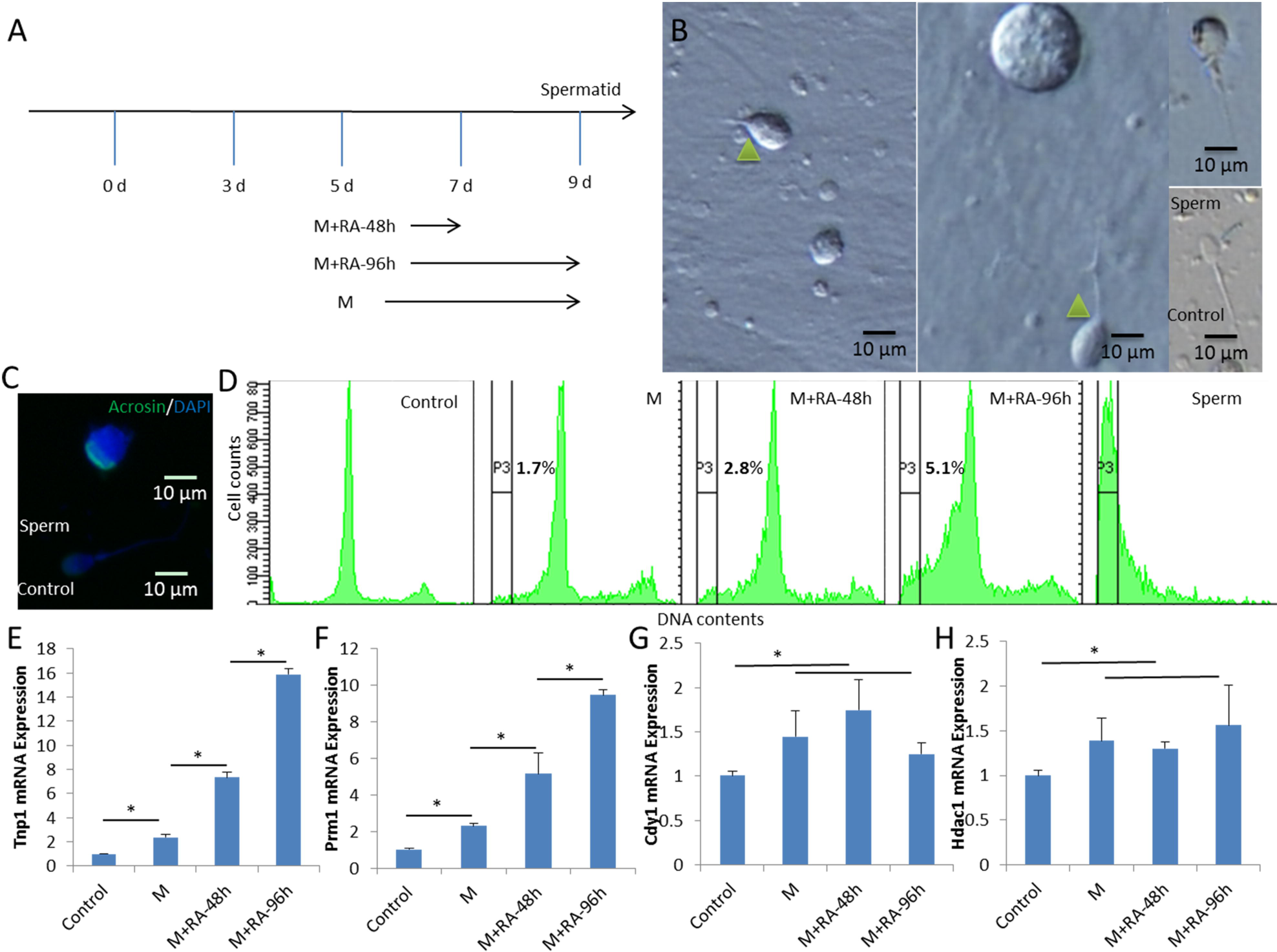
Functional haploid spermatozoa were obtained from *in vitro* culture. A) A schematic illustration of the differentiation process in the present study. B) Representative micrographs of a spermatid with a single flagellum isolated from *in vitro* culture and adult sperm used as a control. C) Haploid cells expressed the mature sperm protein acrosin (green), cell nuclei were stained with DAPI (upper panel), and adult sperm were used as a control (lower panel). D) DNA content of suspended cultured cells was examined by flow cytometry. Control is the group without induction (SSCs on day 5 of incubation), M is the group that was induced to differentiate with basic medium, and M+RA is the group with RA treatment. Adult sperm cells were used as a positive control. P3 marks the haploid peaks. E) and F) Expression patterns of post-meiotic genes (Prm1 and Tnp1). G) and H) Histone acetylation modified enzyme gene Cdyl (and Hdac1) expression. Data are expressed as mean ± SEM. \**P* < 0.05.

### Retinoic acid up-regulated the expression of CREB in porcine SSCs in vitro

The expression of RARγ mRNA was significantly lower in the RAR inhibitor BMS493 group than the M+RA-96h group after 9 days of induction. The content of CREB protein was significantly higher in the M+RA-96h group than that in other groups, and the content of CREB in the RAR inhibitor BMS493 group was lower than that of M+RA-96h group (Figure 4A and B). The addition of RA promoted the expression of anti-apoptotic Bcl2 mRNA, but the addition of BMS493 inhibited this elevation (Figure 4C). These results suggest that RA promoted the post-meiotic germ cell expression of CREB through its specific receptors (Figure 4D).

**Figure 4:**
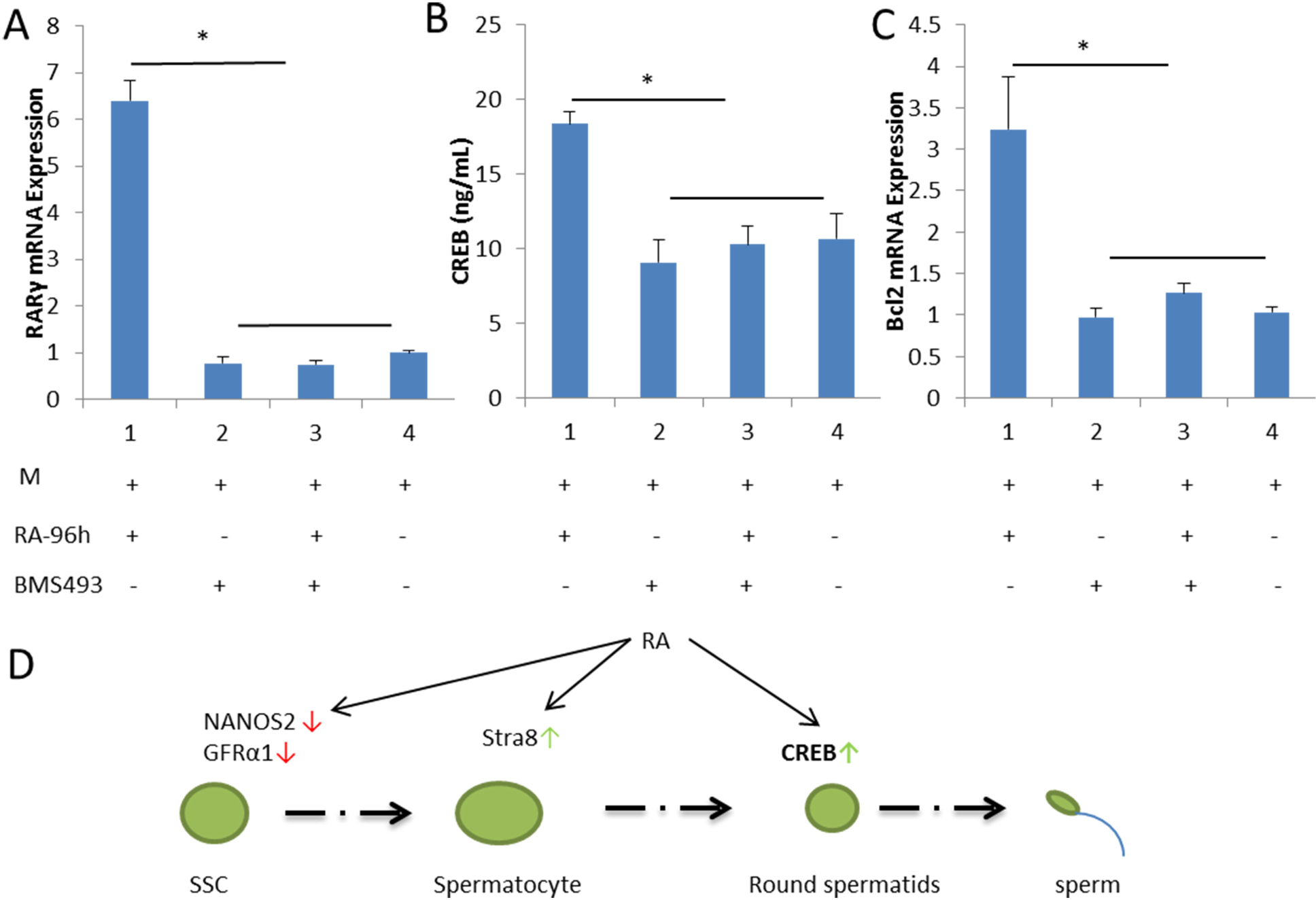
Retinoic acid (RA) up-regulated the expression of cAMP responsive-element binding protein (CREB) in the porcine SSC in vitro differentiation system. A) Real-time PCR analysis of RARγ mRNA levels in the in vitro system on day 9 of incubation. B) CREB levels by ELISA. C) Real-time PCR analysis of Bcl2 mRNA levels in the in vitro system on day 9 of incubation. M is the group that was induced to differentiate with basic medium, and M+RA is the group with RA treatment. Data are expressed as mean ± SEM; \**P* < 0.05. D) RA regulates the SSC differentiation pathway.

### The cultured porcine haploid spermatozoa exhibit developmental potential

Cultured pig round spermatids were injected into metaphase II-stage oocytes (Fig. 5A and B). Injected oocytes formed double-pronuclear reconstructed embryos, as shown by orcein staining (Fig. 5C), and further developed to cleavage and blastocyst stages (Fig. 5D). The rate of blastocyst injection (14.62 ± 3.12%) was significantly lower than that of single sperm injection group (24.60 ± 2.75%) (*P* < 0.05), but had no significant difference with the in vivo round sperm group (16.36 ± 2.25) (Table 3). These findings indicate that the culture-derived pig spermatid with single flagellum had developmental potential in vitro.

**Table 3.**
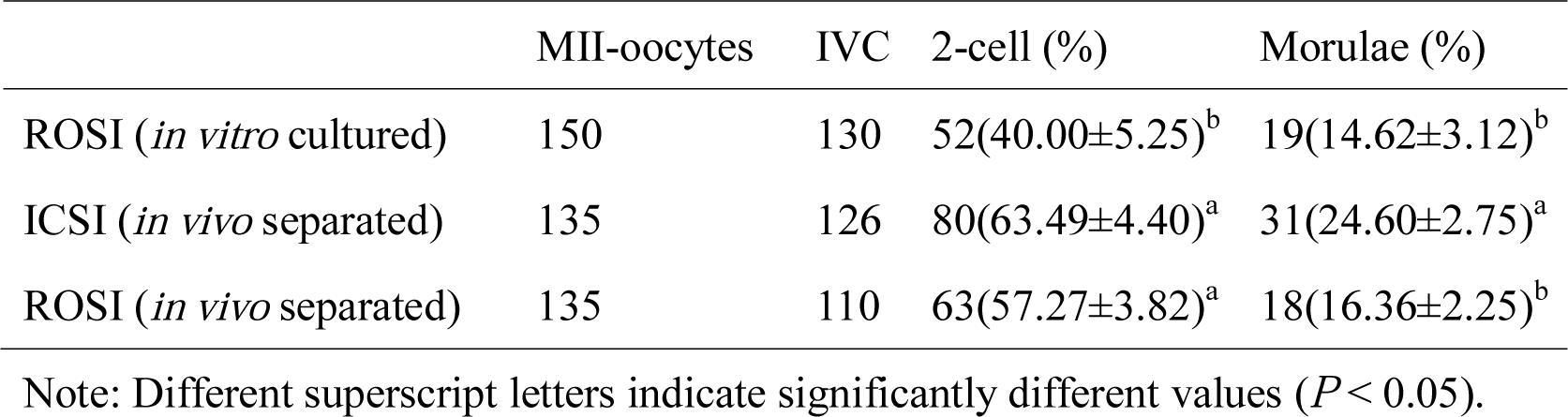
Embryo development of pig oocytes after intracytoplasmic spermatozoa injection (ICSI) or round spermatid injection (ROSI).

**Figure 5:**
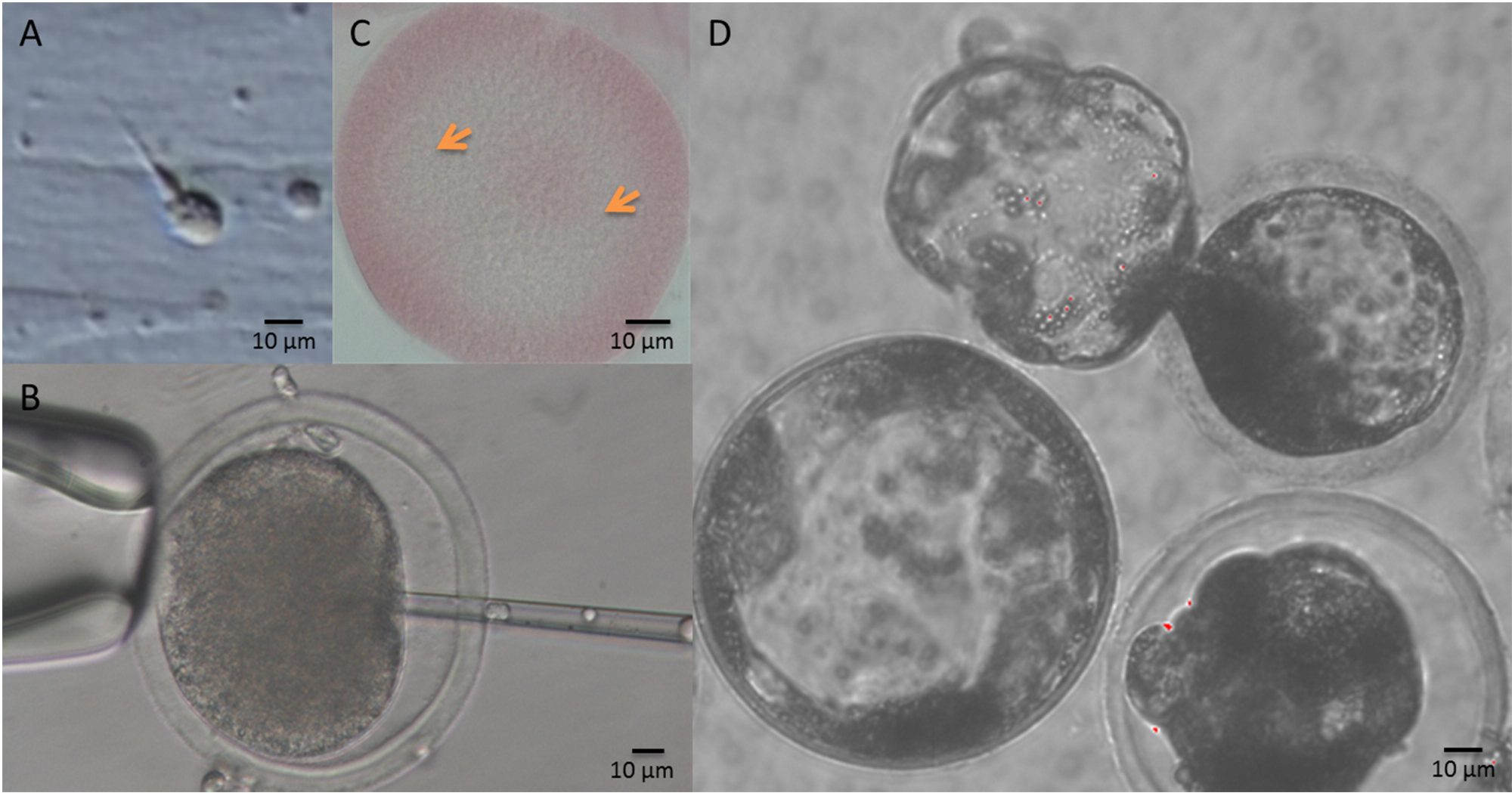
Functional haploid spermatozoa obtained from in vitro differentiation. A) Single tail spermatid obtained from in vitro differentiation. B) Spermatid intracytoplasmic injection into an oocyte. C) Nuclear reconstructed embryos. D) Reconstructed embryos developed to the blastocyst stage.

## Discussion

In this study, porcine SSCs were successfully induced to differentiate into functional haploid spermatozoa in vitro. By adding RA, the differentiation efficiency of haploid cells was enhanced. The RA found in testes is mainly derived from intratesticular synthesis, and testes express a variety of transporters and enzymes related to the synthesis and metabolism of RA. Retinoic acid can be degraded by CYP26B1, which is localized in perivascular myocyte-like cells and regulates RA expression levels in the seminiferous epithelium [35]. Disruption of key enzyme genes in RA synthesis, such as Rdh10 or Aldh1a1–3, leads to RA-deficient mouse testes with spermatogenic arrest at the stage of undifferentiated spermatogonia [36, 37]. Retinoic acid regulates spermatogonial differentiation, spermatocyte meiosis and later stages of spermatogenesis [38]. Retinoic acid triggers spermatogonial differentiation via direct or indirect downregulation of the zinc finger PLZF protein [39], which maintains SSCs in an undifferentiated state. In addition, RA directly actives the phosphorylation of Kit, which regulates the synthesis of DNA in mitotic spermatogonia and the initiation of meiosis via MAPK and PI3K signaling pathways [40]. In mice, lacking the RA target gene Stra8, undifferentiated spermatogonia accumulated in unusually high numbers as early as 10 days after birth, whereas differentiating spermatogonia were depleted [41]. The RNA binding protein NANOS2 can silence genes involved in spermatogonial cell differentiation and meiotic entry, such as stra8, and it is required to maintain the function and survival of undifferentiated spermatogonia [42]. In addition, RA induced undifferentiated spermatogonial cells to form differentiated spermatogonial cells in vitro [43]. The current results from the in vitro differentiation of porcine SSCs showed that RA downregulated the expression of NANOS2, GFRα1 and PLZF in spermatogonia, but promoted the expression of Stra8 in meiotic spermatocytes, and also downregulated the expression of CYP26B1, and promoted the initiation of meiosis.

The formation of mature sperm is associated with RA. In RARα knockout mice, the first wave of spermatogenesis is blocked at step 8 spermatids, but can be rescued by the specific overexpression of RARα in round spermatids [44]. The RAR antagonist BMS-189453 blocks mouse spermatogenesis [45]. Cyclic AMP plays a role in the activation of postmeiotic genes, such as Prm and Tnp, and many genes involved in meiosis include CREB sequence [46]. CREB is an important transcription factor that is differentially regulated in various cell types. The cAMP responsive element modulator (CREM) is an important transcriptional activator during spermatogenesis, especially in postmeiotic germ cells [47]. Inactivation of CREM in mice resulted in an increased rate of apoptosis in the round spermatozoon stage, and the simultaneous expression of apoptosis-related genes [48]. Retinoic acid rapidly activates CREB without using RARs in normal human tracheobronchial epithelial cells. Retinoic acid rapidly activates protein kinase C and transmits an activation signal to phosphorylate nuclear CREB via the Ras/ERK/Rsk pathway, thereby increasing its transactivation activity [49]. Shan et al. [50] found that active CREB protein was increased after treatment with 5 μM RA during the differentiation/formation of the embryoid body. In the *in vitro* induction culture system for porcine spermatogenic cells, RA significantly increased the differentiation rate of haploid germ cells. Retinoic acid can promote the expression of CREB in post-meiotic spermatogenic cells and promote the rate of sperm formation in vitro. This study also revealed that elevated expression of CREB and up-regulated expression of Bcl2 was associated with decreased apoptosis of the cultured porcine reproductive cells in vitro.

## Conclusions

In this study, we successfully used the in vitro culture model of porcine small seminiferous tubule segments to induce SSCs to differentiate into functional single-tail haploid spermatozoa with the potential of further development. When spermatogenic cells in the in vitro culture system were treated with RA, the expression of Stra8 and CREB was up-regulated, likely enhancing the efficiency of producing haploid cells. Through RAR, RA promotes CREB expression, which supports more efficient spermatid differentiation and sperm production.

## Abbreviations

SSCs: Spermatogonial stem cells
RA: Retinoic Acid
GDNF: Glial Cell-Derived Neurotrophic Factor
GFRα1: GDNF Family Receptor Alpha 1
Stra8: Retinoic Acid Gene 8
FSH: Follicle Stimulating Hormone
LH: Luteinizing Hormone
RAR: Retinoic Acid Receptor
CYP26B1: Cytochrome P450 family 26 enzymes B1

## Declarations Acknowledgments

We thank Charles Allan, PhD, from Liwen Bianji, Edanz Editing China (www.liwenbianji.cn/ac), for editing the English text of a draft of this manuscript.

## Author Contributions

Conceived and designed the experiments: Zheng-Xing Lian and Yi-Xun Liu. Performed the experiments: Shou-Long Deng, De-Ping Han and Kun Yu. Analyzed the data: Kun Yu. Contributed reagents/materials/analysis tools: Su-Tian Wang, De-Ping Han and Han-Yu Wu. Wrote the paper: Bao-Lu Zhang, Yi Zhang and Kun Yu.

## Ethics approval and consent to participate

Piglet surgical biopsy was performed at the experimental station of the China Agricultural University, and carried out in strict accordance with the protocol approved by the Animal Welfare Committee of the China Agricultural University.

## Availability of data and materials

The authors confirm that all data generated or analyzed during this study are available.

## Consent for publication

Not applicable.

## Competing financial interests

The authors declare that they have no competing interests.

**Supplementary movie. A spermatid with a single flagellum from *in vitro* culture.**

